# Multiomics Integration Reveals a Metabolic Myopathy in Cardiometabolic HFpEF

**DOI:** 10.64898/2025.12.15.694522

**Authors:** Timothy D. Allerton, Daniel Arabie, Kade Malone, Abhishek Pandit, Mahmoud H. Elbatreek, Zhen Li, Robert C. Noland, Jake Doiron, Mike Kinter, Brooke Loveland, Pravalika Javvadi, Kirti Agrawal, Manas Gartia, Praveen Guttula, Traci T. Goodchild, Sanjiv J. Shah, Sujoy Ghosh, David J. Lefer

## Abstract

**Background:** Skeletal muscle dysfunction is a major peripheral determinant of exercise intolerance and physical disability in heart failure with preserved ejection fraction (HFpEF). Metabolic and mitochondrial dysfunction are considered to be key components of skeletal muscle dysfunction, but comprehensive profiling of metabolic pathways has not been conducted. Elucidation of dysregulated metabolic pathways is essential to determine viable targets for the treatment of exercise intolerance in HFpEF.

**Methods:** Male ZSF1 Obese rats (HFpEF) and Wistar Kyoto (WKY) lean normotensive controls were studied at 26 weeks of age. Gastrocnemius was subjected to bulk RNA-seq, proteomics, metabolomics, and lipidomics analysis. The R package *limma* was used to determine differential expression in all omics layers (absolute fold-change>1.5, FDR0.05, unless otherwise indicated). Additional targeted plasma and skeletal muscle (soleus and EDL) metabolomics and lipidomics were performed on HFpEF and control rats.

**Results:** Pathway level analysis for RNA seq and proteomics revealed significant downregulation of oxidative phosphorylation (NES -2.1, p<0.005), electron transport chain (NES -2.0, p<0.005), and TCA cycle (-1.8, p<0.05). The most upregulated pathways were PPAR signaling (NES 2.2, p<0.0001), tryptophan metabolism (NES 1.8, P<0.005), and amino acid oxidation (NES 1.8, p<0.005) pathways. Metabolomics revealed an accumulation of TCA cycle intermediate, isocitrate, and phosphate reduction. Branched-chain amino acids were significantly increased, whereas amino acids related to tryptophan metabolism were reduced and shifted towards increased serotonin accumulation. Phospholipid species were differentially regulated with increased palmitoylated phosphatidylcholines but reduced arachidonoyl-PC species. Phosphatidylethanolamines (PE) species (16:0/16:1-18:0/18:2) were increased.

**Conclusion:** Our multiomics analysis of skeletal muscle in HFpEF revealed severe mitochondrial dysfunction that was characterized by reduced complex I and II activity. Mitochondrial and peroxisomal lipid overload results in a shift in membrane phospholipid accumulation and composition. Reduced BCAA oxidation and dysregulation of tryptophan metabolism are key features of amino acid metabolism that reduce anaplerosis and promote the accumulation of toxic metabolites. Comparative analysis of other skeletal muscle disorders suggests that an acquired metabolic myopathy exists in cardiometabolic HFpEF.

## INTRODUCTION

Heart failure with preserved ejection fraction (HFpEF) is now the predominant form of heart failure in the United States, afflicting approximately 3 million people ^1^. The global prevalence of HFpEF has risen sharply in recent years, with current estimates showing it now accounts for approximately 20–50% of all heart failure cases worldwide and has contributed to the doubling of total heart failure prevalence to over 56 million people since the 1990s^2^. The long-term prognosis for patients with HFpEF is very poor, and effective treatments are extremely limited. The 5-year mortality rate after diagnosis of HFpEF is 50% and increases to 75% after HFpEF hospitalization^1,3^. Chronic exercise intolerance is a prime symptom of HFpEF and a sensitive predictor of mortality, hospitalization, and poor quality of life ^4–6^. The etiology of exercise intolerance and reduced functional capacity is multifactorial and involves a combination of both cardiac and peripheral factors (i.e., skeletal muscle).

HFpEF is a disease with high phenotypic diversity. Obesity is a major risk factor for the development of HFpEF and defines the cardiometabolic HFpEF phenotype ^7,8^. Patients with cardiometabolic HFpEF have more severe exercise intolerance when compared to other HFpEF phenotypes, largely due to a disproportionately greater degree of skeletal muscle dysregulation ^9–14^. Numerous sites of dysregulation have been noted in the skeletal muscle of HFpEF patients. Early reports described the loss of type I oxidative fibers, reduced capillary density, and adipose tissue infiltration as key pathophysiology features of skeletal muscle in HFpEF patients ^15,16^. Subsequent reports also established that patients with HFpEF experience reduced contraction-initiated skeletal muscle perfusion ^10,12,17–19^. Importantly, these studies established that these deficits were highly associated with the degree of exercise intolerance, independent of changes in central hemodynamics. Muscle weakness (i.e, reduced muscle force generation and fatigue resistance) has also been reported in HFpEF that either rivals or exceeds what is experienced in heart failure with reduced ejection fraction (HFrEF)^13,20^. Collectively, these seminal studies have identified key aspects of skeletal muscle dysfunction that contribute to reduced exercise capacity and muscle strength in HFpEF.

Mitochondrial dysfunction is also implicated as a causal factor limiting exercise capacity in HFpEF. Multiple reports have demonstrated reduced skeletal muscle mitochondrial respiration in patients with HFpEF^13,19,21^. Impaired mitochondrial respiration in the vastus lateralis was highly correlated with peak oxygen consumption (VO_2peak_), 6-minute walk distance, and leg strength^21^. These reports, and others, have established the premise that skeletal muscle bioenergetics is a critical regulator of exercise intolerance in HFpEF^22^. Despite these hallmark studies, the landscape of skeletal muscle dysregulation remains poorly understood. Additionally, many of these studies have characterized non-obese HFpEF patients who are phenotypically distinct from the cardiometabolic HFpEF phenogroup. Recent reports using a multiomics approach have expanded our knowledge of cardiac metabolic metabolism in HFpEF^23,24^. The purpose of the current investigation was to provide a comprehensive analysis of molecular transducers of skeletal muscle dysfunction in a translational model of cardiometabolic HFpEF. We used a multiomics approach (transcriptomics, proteomics, lipidomics, and metabolomics) to elucidate key metabolic pathways in the ZSF1 obese rat model of HFpEF. Our analysis revealed a downregulation of gene expression for the TCA cycle, electron transport, and oxidative phosphorylation. While mitochondrial and peroxisomal β-oxidation are increased, the accumulation of specific metabolic indicates this pathway is incomplete in these organelles. Metabolomics and lipidomics confirm that the overload of these bioenergetic pathways promotes the generation of rare toxic lipid species. We are also the first to report that significant remodeling of membrane phospholipids and the generation of inflammatory lysophospholipids in HFpEF are key metabolic features in HFpEF.

## METHODS

### Animal Models

Male ZSF1 obese (n=8) and Wistar Kyoto (WKY) rats (n=8) were purchased (Charles River Laboratories, Wilmington, MA, USA) at 8 weeks of age and allowed to acclimate for 2 weeks. Male ZSF1 lean rats (n=5) were also used as an additional comparison group and phenotyped identically. Rats were housed at LSUHSC in a temperature-controlled controlled and a 12-hour light/dark cycle. All studies were IACUC (Institutional Animal Care and Use Committee) approved and received care in LSUHSC animal care according to AALAC guidelines. ***Exercise Treadmill Capacity***

Exercise capacity in rats was assessed using an IITC Life Science 800 Series treadmill (Woodland Hills, CA), specifically designed for rodents. Before testing, all animals underwent a 5-minute acclimation period on the treadmill. Rats were acclimated by a warm-up at 6 m/min, with speed gradually increased by 1.5 m/min to a final 12 m/min, maintained for one additional minute. After the 5-minute warm-up, rats performed at 12 m/min on a flat plane until exhaustion. Exhaustion was defined as failure to run for more than 5 seconds or inability to reach the front of the treadmill for 20 seconds. Exercise performance was quantified as both distance and work (kg·m), using body weight measured immediately before testing.

### Skeletal Muscle Grip Force

Forelimb grip strength in rats was assessed at 26 weeks of age using a grip strength meter (GSM, Columbus Instruments, US). The forelimb pull bar was attached to the disinfected GSM. The GSM was activated in Peak Tension (T-PK) Mode and tared to zero. Each rat was held by its tail and lowered towards the pull bar until it grasped it. The rat was then pulled backward in a straight horizontal line until it released its grip. The peak force displayed on the GSM was recorded, and the device was tared again before testing the next animal. Each rat underwent this procedure five times, with several minutes of rest provided between each trial.

### Transthoracic Echocardiography

Echocardiography was performed using a Vevo 2100 system (VisualSonics, Toronto, Canada) according to previously reported methods^25^. Briefly, rats were shaved 18 h before the experiment. Rats were placed on a heated (37 °C) platform and anesthetized with 3% isoflurane, maintained at 1–3% during the procedure. Target heart rates were 300–350 bpm for both systolic and diastolic measures. LVEF was measured from M-mode images in the parasternal short-axis view, and E, A, and E′ velocities were obtained from the apical four-chamber view. Respiratory rate was continuously monitored to ensure welfare and physiologic data quality. All animals recovered uneventfully. The data represent the average of three consecutive measurements per parameter.

### Hemodynamics

Rats were anesthetized (3.0 % isoflurane) until fully unresponsive. Next, the right common carotid artery was surgically isolated and exposed. Isoflurane was reduced to 1%, and a 1.6-Fr (rat) high-fidelity pressure catheter (Transonic, NY, USA) was inserted to record systemic blood pressures. The catheter was then advanced into the left ventricle to measure left ventricular end-diastolic pressure (LVEDP). Respiratory rate was continuously monitored to ensure welfare and measurement accuracy. After hemodynamic assessment, isoflurane was increased to 5% for blood collection via direct cardiac puncture, and animals were euthanized by cardiectomy under anesthesia.

### Histology

Skeletal muscle sections (gastrocnemius) were fixed in formalin and processed according to standard methods. Sections were sliced at stained with hematoxylin and eosin, picosirius red with fast green as a counterstain. Perilipin IHC staining was accomplished using a Leica Bond RXm platform using a modified Leica IHC protocol and utilizing a Bond Polymer Refine Detection kit (Leica Biosystems, DS9800) as the preferred detection system. Briefly, each slide underwent onboard ‘bake and dewax’ followed by heat induced antigen retrieval for 20 minutes using Leica ER2 solution (Leica Biosystems, AR9640) and were processed using a modified “Protocol F” that included 5 minutes of Leica peroxide block, 30 minutes incubation with rabbit anti-perilipin-1 antibody (Cell Signaling Technology, 9349; 1:1000), incubation with Leica’s goat anti-rabbit HRP polymer for 8 minutes, and development with Leica’s mixed DAB Refine for 10 minutes. Slides were counterstained with hematoxylin and coverslipped using a Leica ST5020 autostainer and CV5030 coverslipper. Slides were dewaxed and subjected to heat-induced antigen retrieval for 20 minutes @ 100 °C using a pressure cooker and pH 6.0 citric acid-based unmasking solution (Vector Laboratories, H3300). After cooling for approximately 20 minutes, the slides were washed in TBST (TBS + 0.1% Tween-20) before being subjected to photobleaching for 1 hour between two 1200W LED grow lights while immersed in 4 ml of photobleach solution (4.5% (w/v) H2O2, and 20mM NaOH in PBS). Slides were then rinsed in TBST and incubated overnight at 4C with biotinylated Griffonia Simplicifolia lectin 1 isolectin B4 (Vector Laboratories, B1205; 1:30) in primary antibody diluent (Leica Biosystems, AR9352). After washing the slides three times for 5 minutes each in TBST, they were incubated with Cy3-conjugated streptavidin (Jackson ImmunoResearch, 016-160-084; 1:500) in TBST for 1 hour and then incubated overnight at 4C with rabbit anti-laminin antibody (abcam, ab11575; 1:000). The slides were washed three times for five minutes in TBST and incubated for 2 hours with a donkey anti-rabbit secondary antibody (Jackson Immunoresearch, 711-605-152; 1:300). Samples were washed once more in TBST, counterstained with Hoechst, and mounted with Aqua-Poly/Mount (Polysciences, 18606).

### Mitochondrial Complex I & II Activity

Frozen tibialis anterior muscles were pulverized under liquid N_2_, and approximately 200 mg of powdered tissue was suspended in 5 volumes of ice-cold assay buffer [10 mM Tris-SO_4_ (pH 7.8 at 30°C), 250 mM sucrose, 1 mM EDTA, and 1mg/mL BSA]. The suspended tissue was then homogenized by performing 15 passes at 1000 RPM with a Potter-Elvehjem tissue homogenizer with a motor-driven PTFE pestle. Insoluble debris was subsequently removed by centrifugation at 1000 xg for 10 min at 4°C. The supernatants were retained, flash-frozen in liquid N_2_, and stored at −80°C. Muscle protein concentrations were determined by using the Pierce™ 660 nm Protein Assay (Thermo Fisher Scientific, Rockford, IL, USA) according to the manufacturer’s protocol. Prior to performing enzyme assays, samples were thawed on ice and diluted to 1.5 mg·mL^−1^ of muscle protein in assay buffer (described above). Complex I activity of tissue homogenates was measured according to the method previously described by Burger et al. (2023) with minor modifications^26^. The 200 µL reaction mixtures were prepared in microplate wells containing 10 mM Tris-SO_4_ (pH 7.8 at 30°C), 250 mM sucrose, 1 mM EDTA, 1 mg·mL^−1^ BSA, 1 µM antimycin A, 200 µM KCN, 100 µM decylubiquinone, 200 µM NADH, 30 µg muscle protein, and either 2 µM rotenone or ethanol (vehicle). Reactions were initiated by the addition of NADH, and the rate of NADH oxidation at 30°C was monitored spectrophotometrically for 5 min at 340-380 nm (ε_340-380_ = 4.81 mM^−1^·cm^−1^;)^27,28^ using a Spectramax 384 Plus microplate reader (Molecular Devices; San Jose, CA, USA). Duplicate assays were performed on each sample. Parallel reactions were performed in the presence of the Complex I inhibitor rotenone, and the rotenone-insensitive rate was subtracted from the uninhibited rate to determine the NADH:decylubiquione oxidoreductase activity. Assays of Complex II activity were performed using a method adapted from a previously published protocol^29^. Duplicate assays of each sample were performed in microplate wells with a final volume of 200 µL containing 10 mM Tris-SO_4_ (pH 7.8 at 30°C), 250 mM sucrose, 1 mM EDTA, 1 mg·mL^−1^ BSA, 1 µM antimycin A, 200 µM KCN,100 µM 2,6-dichlorophenolindophenol (DCPIP), 5 mM succinate, 100 µM decylubiquinone, 30 µg muscle protein, and either 10 mM malonate or distilled H_2_O (vehicle). Samples were pre-incubated in the reaction mixture described above without decylubiquinone at 30°C for 10 min to allow succinate-dependent activation of Complex II, and reactions were initiated by the addition of decylubiquinone^29^. The reduction of DPCIP was monitored by measuring the decrease in Abs_600_ at 30°C for 5 min. The rate of DCPIP reduction was determined by using the DCPIP extinction coefficient (ε_600_ = 19.1 mM^−1^·cm^−1^). The Complex II inhibitor malonate was included in parallel reactions to determine the background rate of DCIP reduction, which was subtracted from the uninhibited rates to determine the specific activity of succinate dehydrogenase.

### Immunoblotting

Tissue was lyophilized with a mortar and pestle and then homogenized in 700 Pierce™ IP Lysis (Thermo Scientific™, 87787) supplemented with a protease inhibitor (Millipore Sigma™, 539131) and phosphatase inhibitor (Thermo Scientific™, 1862495) cocktail. These lysates were then centrifuged (16.000 xg, 15 minutes, 4 °C) and supernatants were collected. Protein concentration was quantified using the Pierce™ 660 nm Protein Assay Reagent (Thermo Scientific™, 1861426). Concentrations were normalized and dispensed in 20 μG input aliquots before Western blot analysis. Each sample was prepared with the addition of Laemmli buffer (BIO-RAD, 1610747) and 10% β-mercaptoethanol (Sigma-Aldrich, M6250). Samples were separated on 10% SDS-PAGE and transferred to a nitrocellulose membrane (Thermo Scientific™, 88018). After transfer, the blots were stained with No-Stain™ Protein Labeling Reagent (Thermo Scientific™, A44449) for total protein normalization in downstream analysis. The membranes were then blocked with EveryBlot Blocking Buffer (BIO-RAD, 12010020) and probed with anti-BCKDH-E1-α (Cell Signaling, 90198S) and anti-P-BCKDH-E1-α (Cell Signaling, 40368S). Blots were then probed with anti-rabbit secondary antibody (Cell Signaling, 7074P2) and visualized using SuperSignal™ West Pico PLUS Chemiluminescent Substrate (Thermo Scientific™, 34580). Quantification and normalization of Western blots were performed using the iBright Analysis Software (Thermo Scientific™, Waltham, MA).

### RNA-Seq

RNA was isolated from skeletal muscle (gastrocnemius, soleus, and EDL) according to established methods ^30,31^. Library construction was performed using Lexogen Quant-Seq with 50 bp forward reads. Sequencing and initial data processing were performed on an Illumina NextSeq instrument according to previously established methods ^32,33^. Sequencing reads were quality-checked via the RseQC package (version 5.0.3)^34^, adapter trimmed via Cutadapt^35^, mapped to the rat reference genome (Rnor_6.0 release 110) via STAR (version 2.7.11b)^36^, and converted to raw counts via the HTSeq package (version 2.0.5) ^37^. Raw counts were normalized via the TMM-normalization method in the edgeR package (v. 3.38.4) ^38^. Principal component analysis was conducted via the princomp function in R for outlier detection. Differential gene expression analysis was performed via the ‘trend’ function in the limma package (version 3.56.1) ^39^, and genes with absolute fold-change <1.5 and FDR < 0.05 were considered as differentially expressed. Gene-set enrichment analysis was performed via the Fast Gene Set Enrichment Analysis (fgsea) (v. 1.30.0) package ^40^, based on gene set permutations on a pre-ranked gene list to identify pathway enrichment. Three pathway databases were queried from the Molecular Signatures Database (MSigDB)^41^: Hallmark, Wikipathways, and Kyoto Encyclopedia of Genes and Genomes (KEGG; https://www.genome.jp/kegg/pathway.html). Top-scoring pathways were defined as those with adjusted P values for enrichment <0.05 (unless reported otherwise). Ingenuity Pathway Analysis Software (Qiagen, Redwood City, CA, www.qiagen.com/ingenuity) was used to determine canonical pathways for comparative analysis, as previously established ^42^.

### Proteomics

A total of 50 µg of homogenized gastrocnemius was used for proteomics analysis. Each sample was mixed with 150 µL 1% SDS, 20uL 10% SDS, and 20 µL of BSA internal standard (1ug). The samples were mixed, heated at 70°C, and precipitated with acetone overnight. The precipitate was reconstituted in 50 µL of Laemmli sample buffer, and 20 µL was run 1.5cm into an SDS-Page gel. Each 1.5cm lane was cut as a sample, cut into smaller pieces, washed, reduced, alkylated, and digested with 1 µg trypsin overnight at room temperature. Peptides were extracted from the gel in 50% acetonitrile, the extracts taken to dryness by Speedvac, and reconstituted in 200 µL 1% acetic acid for analysis. Analysis was conducted using Orbitrap Exploris^TM^ 480 Mass Spectrometer in DIA mode with a 10m/z window working from m/z 350 to 950. The Orbitrap was operated at a resolution of 15,000. A full scan spectrum at a resolution of 30,000 was acquired each cycle. Data were analyzed using the program DIA-NN. This program uses neural network methods to analyze the DIA datasets versus the respective Uniprot proteome database from EMBL.org. Data are reported as units pmol/100ug total protein and are based on normalization to the BSA internal standard added to all samples. As the proteomics data contained missing values not at random (MNAR), we performed imputation of the missing values via the QRILC method through the DEP R package (v. 1.28.0), after removing proteins with >50% missing data^43^. Imputed data was subjected to an additional round of variance stabilizing normalization via vsn^44^. Principal components analysis of the normalized proteomics data via the princomp function in R detected no outliers. Differential proteomic analysis was carried out via the trend function in the limma package (v 3.60.6)^39^. Pathway over-representation analysis was performed via Enrichr^45^ through interrogation of biological pathways from the Wikipathways database ^46^.

### Global Metabolomics

Gastrocnemius samples were processed using the automated Hamilton MicroLab STAR® system. Recovery standards were added before extraction for quality control. Proteins were precipitated with methanol under vigorous shaking for 2 minutes, followed by centrifugation, to remove protein, release small molecules bound within protein matrices, and recover chemically diverse metabolites. The resulting extract was divided into multiple fractions: two for analysis by distinct reverse-phase (RP) UPLC–MS/MS methods using positive-ion electrospray ionization (ESI); one for RP/UPLC–MS/MS using negative-ion ESI; one for HILIC/UPLC–MS/MS using negative-ion ESI; with additional fractions retained as backup. All analyses were performed on a Waters ACQUITY UPLC system coupled to a Thermo Scientific Q-Exactive high-resolution/accurate-mass spectrometer equipped with a HESI-II source and an Orbitrap mass analyzer operated at 35,000 mass resolution. Dried extracts were reconstituted in solvents specific to each of the four analytical methods, each containing a set of internal standards to ensure consistent injection and chromatography. One aliquot was analyzed under acidic positive-ion conditions optimized for hydrophilic compounds (PosEarly). Samples were gradient-eluted from a Waters UPLC BEH C18 column (2.1 × 100 mm, 1.7 µm) using water and methanol containing 0.05% PFPA and 0.1% FA. A second aliquot was analyzed under similar acidic positive-ion conditions but optimized for hydrophobic analytes (PosLate). These extracts were gradient-eluted from the same C18 column using methanol, acetonitrile, water, 0.05% PFPA, and 0.01% FA at a higher overall organic content. A third aliquot was analyzed using basic negative-ion conditions (Neg) on a dedicated C18 column, with gradient elution using methanol and water containing 6.5 mM ammonium bicarbonate at pH 8. The fourth aliquot was analyzed by negative-ionization HILIC (HILIC) using a Waters UPLC BEH Amide column (2.1 × 150 mm, 1.7 µm) eluted with a water–acetonitrile gradient containing 10 mM ammonium formate at pH 10.8. MS acquisition alternated between full MS and data-dependent MSⁿ scans with dynamic exclusion. Scan ranges varied slightly between methods but generally spanned 70–1000 m/z. Raw data files were archived and processed as described below. Before storage, samples were briefly dried on a TurboVap® (Zymark) to remove residual organic solvent. Extracts were then held overnight under nitrogen before analytical preparation. Metabolite data was imputed and batch corrected at Metabolon, followed by median-based normalization in Metaboanalyst^47^. Normalized data was log2 transformed and subjected to differential metabolite expression analysis via the trend function in limma (v. 3.60.6)^39^.

### Targeted Metabolomics

Plasma and skeletal muscle samples were processed using a 1:5 ratio of ice-cold HPLC-grade isopropanol. A targeted metabolomics analysis was performed according to the manufacturer’s instructions using the Biocrates MxP Quant 500 kit (Biocrates, Innsbruck, Austria), which enables the quantification of 630 metabolites from 10 μL of human plasma. In brief, metabolite measurements were conducted on a Nexera HPLC system (Shimadzu) coupled with a 6500+ QTRAP mass spectrometer (AB Sciex) employing an electrospray ionization source, following established protocols^48^. Mass spectrometry data were processed using Sciex Analyst software and subsequently analyzed in Biocrates MetIDQ software. All measurements were normalized using internal quality control standards.

### Statistical Analysis

The normality of data distribution was determined using D’Agostino & Pearson and Shapiro-Wilk tests. Where appropriate, two groups were compared using an unpaired *t*-test for normally distributed data; a Welch’s t-test was used for data that was not normally distributed. The difference in the number of rats in the presented data reflects a subset of tissue used for certain analysis due to the limitation of tissue. All data are presented as mean ± standard deviation. Statistical significance was determined as P<0.05. Statistical analysis was performed using Prism 10 software (GraphPad, San Diego, CA).

## RESULTS

### Metabolic Pathways Dominate the Omics Signature of Skeletal Muscle in HFpEF

ZSF1 obese (HFpEF) and WKY rats were aged to 26 weeks of age (Figure 1A). HFpEF rats had increased body weight (Figure 1B), exercise intolerance (Figure 1C-D), and skeletal muscle weakness (Figure 1E). We also confirmed the cardiac HFpEF phenotype defined by impaired diastolic function (Supplemental Figure 1A) and increased left ventricular filling pressure (Supplemental Figure 1B). Our histological analysis of skeletal muscle pathology revealed a decrease in muscle fiber cross-sectional area (Figure 1F), increased interstitial (Figure 1G) and perivascular fibrosis (Figure 1H), and reduced capillary density (Figure 1I). We also performed immunohistochemistry using perilipin 1 to demonstrate adipocyte infiltration in the HFpEF skeletal muscle (Figure 1J). Next, we performed bulk RNA sequencing, proteomics, and metabolomics on snap-frozen gastrocnemius samples (Figure 1K). Transcriptomics revealed a large number (1242 genes, 646 decreased & 596 increased) of differentially regulated genes (Figure 1L-M, Supplemental Table 1.1) between HFpEF and control rats. Fewer genes (686 genes) were differentially expressed when comparing ZSF1 Obese to ZSF1 lean rats or ZSF1 lean to WKY (353 genes) (Supplemental Figure 1.2-3). Pathway-level analysis (Figure 1N) showed significant enrichment (natural enrichment score, NES) of metabolic pathways (Wikipathways) in ZSF1 obese (HFpEF) rats. Oxidative phosphorylation (NES=-2.13, Adjusted P<0.0001), electron transport chain (NES=-2.04, Adjusted P<0.0001), and TCA cycle (NES=1.84, Adjusted P=0.03) were all significantly downregulated. Whereas, PPAR signaling (NES=2.27, Adjusted P<0.0001), tryptophan metabolism (NES=1.85, Adjusted P=0.02), and amino acid metabolism (NES=1.79, Adjusted P<0.005) were significantly upregulated pathways. Proteomics detected 1765 proteins, of which 334 were differentially expressed (Figure 1O, Adjusted P<0.05), and the top 50 most differentially regulated are shown in Figure 1P. We confirmed significant enrichment of pathways related to fatty acid oxidation, amino acid metabolism, and the electron transport chain (Figure 1Q). Proteomics also showed enrichment of glutathione and glycogen metabolism, providing further evidence that metabolic pathways are the predominant site of dysfunction in skeletal muscle in HFpEF. The most prominent pathways were altered regardless of the control group (Supplemental Figure 2-3). Untargeted metabolomics revealed metabolites such as 5-aminovalerate, imidazole lactate were upregulated, while X-23171 (unidentified) and 1-margaroyl-2-arachidonoyl-GPC (17:0/20:4) were among the most reduced metabolites in the skeletal muscle (Figure 1R-S). Numerous metabolites related to glucose, glutathione disulfide, and glycation production support oxidative stress and metabolic overload in the HFpEF skeletal muscle (Supplemental Table 2.1). Phospholipid metabolism was highly altered in HFpEF, with 78 lipid species (Supplemental Table 2.1) being differentially regulated.

**Figure 1.**
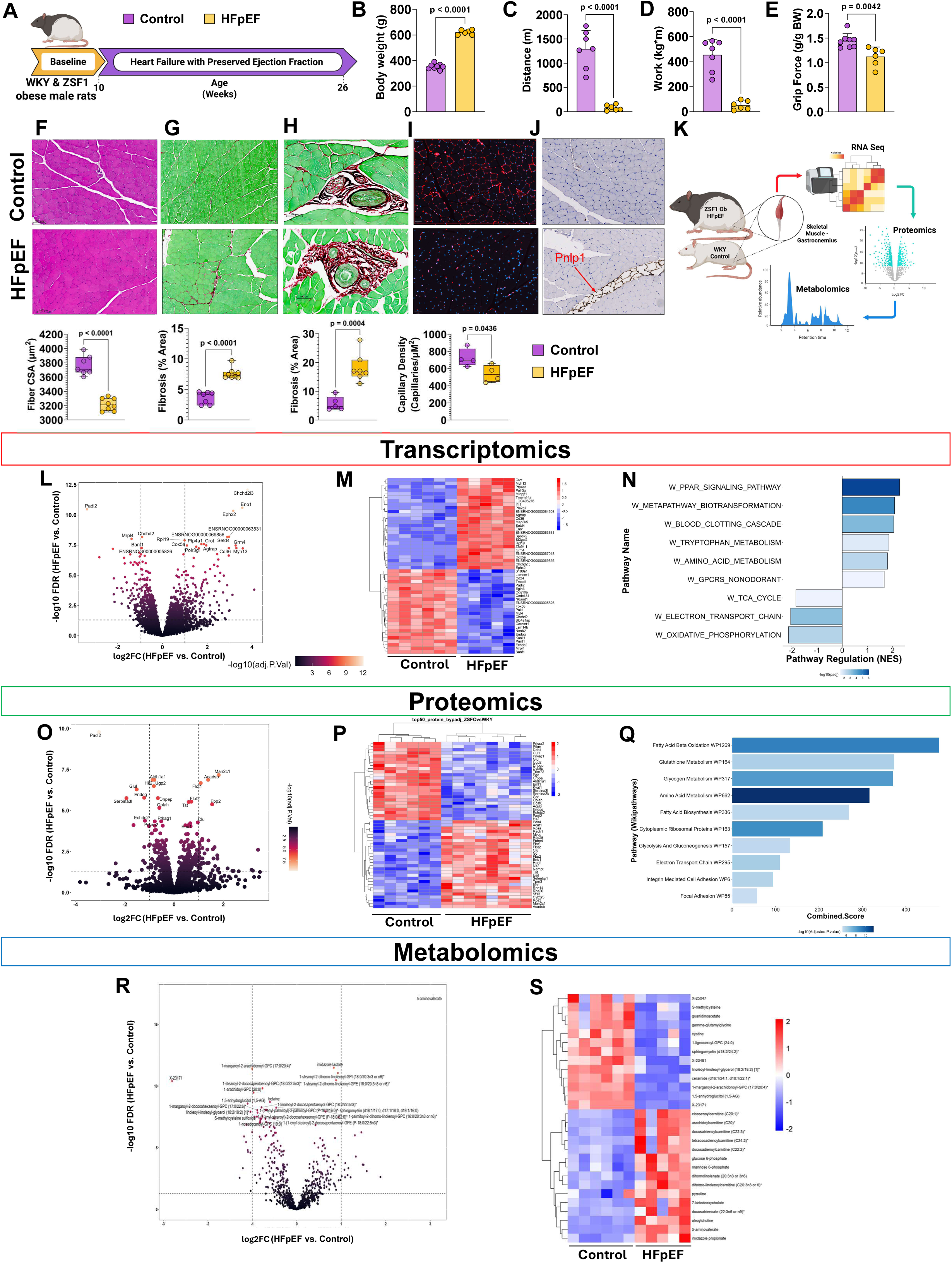
Multiomics of skeletal muscle in ZSF1 Obese Rats Model of Cardiometabolic HFpEF. **(A)** Study design of ZSF1 obese (HFpEF) and WKY (control) rats studied at 26 weeks of age. Gastrocnemius was harvested at the time of sacrifice for downstream transcriptomics, proteomics, and metabolomics analysis. **(B)** Body weight, **(C)** treadmill exercise distance **(D)** work, and **(E)** forelimb grip force production in HFpEF versus control rats at 26 weeks of age. **(F-J)** Histological examples and quantification (below) of **(F)** H&E, **(G-H)** picosirius red, **(I)** IB4-isoselectin, and (J) perilpin 1 immunostaining of gastrocnemius sections. **(K)** Multiomics integration design. **(L)** Volcano plot and **(M)** heatmap for the most differentially regulated gene identified with the bulk RNA-seq approach. **(N)** Wiki pathways for the most enriched pathways in HFpEF versus control rats. **(O)** Volcano plot and **(P)** heatmap for the most differentially regulated proteins from proteomics analysis. **(Q)** Wikipathways for the most enriched pathways in HFpEF versus control rats. **(R)** Volcano plot and **(S)** heatmap of the most differentially regulated metabolites from global metabolomics analysis. All data are mean ± SD. An unpaired *t*-test was used to determine significance for figures B-I. An FDR <0.05 was used to determine significantly altered genes, proteins, and metabolites. n=6-8/group. Significant values are provided within graphs.

Cardiometabolic HFpEF is a heterogeneous disease characterized by obesity, hypertension, and chronic inflammation. However, at the level of the skeletal muscle, there is little context beyond the observation that skeletal muscle demonstrates numerous features of obesity and insulin resistance. We utilized our RNA Seq analysis to conduct a comparative analysis of publicly available datasets (IPA, Qiagen Curated Pathways) to determine similarities between other disease states and cardiometabolic HFpEF. Surprisingly, muscle myopathies and dystrophies were the top-scoring matches in our analysis (Table 1). Polymyositis, facioscapular-humeral muscle dystrophy, Duchenne’s muscle dystrophy, and limb-girdle muscular dystrophy type 2A had similar gene expression patterns the the ZSF1 obese rat. These disorders are associated with chronic inflammation and progressive muscle weakness. Other matching pathways included chronic inflammation of traumatic injury, aging, sarcopenia, and COPD.

**Table 1.** Ingenuity Pathway Analysis Disease Match.

| **Table 1. Ingenuity Pathway Analysis Disease Match** |  |  |  |
| --- | --- | --- | --- |
| **Disease State** | **Species** | **Project Name** | **Z-score** |
| Polymyositis | Human | GSE26852 | 44.8 |
| Aging | Human | GSE8479 | 43.4 |
| Wounds and Injury | Rat | GSE162565 | 42.2 |
| Facioscapulohumeral muscular dystrophy | Human | GSE26852 | 41.2 |
| COPD | Human | GSE100281 | 34.3 |
| Dermatomyositis | Human | GSE26852 | 30.0 |
| Sarcopenia | Rat | GSE146976 | 29.6 |
| Duchenne muscular dystrophy | Human | GSE3307 | 28.2 |
| Limb girdle muscular dystrophy type 2A | Human | GSE3307 | 27.4 |

A multiomic (gene, protein, and metabolite) integration (Figure 2A-B) shows gene expression was mostly downregulated in glycolysis (Figure 2C), except for Eno1. TCA cycle gene expression (Figure 2D) was predominantly downregulated. Figure 2E-H confirms the upregulation of Eno1, and the reduction of Aldoa, PFK, and HK proteins. Glucose, glucose-6-phosphate, and fructose-6-phosphate were significantly accumulated glycolytic metabolites (Figure 2I). Isocitrate was the only increased TCA cycle intermediate (Figure 2J). Succinate was numerically increased (2.16 FC), but did not reach significance (P=0.10, Supplemental Table 2.2). NADH, NADH: NAD^+^, and purine nucleotides such as adenosine 3’-monophosphate (3’-AMP), adenosine 3’,5’-diphosphate, and adenosine 3’,5’-cyclic monophosphate (cAMP) were increased as were also significantly increased in HFpEF versus control skeletal muscle (Figure 2K). TCA intermediates can be exported from the skeletal muscle into the circulation under cellular stress. Therefore, plasma carboxylic acids were measured in the plasma using targeted LC-MS. Results showed significantly elevated aconitic acid (via the aconitase) and lactic acid, with numerical increases that neared significance in hydroxyglutaric acid (via the reduction of α-ketoglutarate) and succinic acid (Figure 2L). Both complex I and II activities were reduced in HFpEF skeletal muscle (tibialis anterior) (Figure 2M). Differential expression analysis of gene expression for mitochondria-associated genes from the MitoCarta (3.0) database (Figure 2N) shows a reduction of the Complex I (NADH: Ubiquonone) assembly genes. Inner membrane and respiratory complexes were particularly enriched mitochondrial cellular components (Figure 2O), altered in HFpEF skeletal muscle.

**Figure 2.**
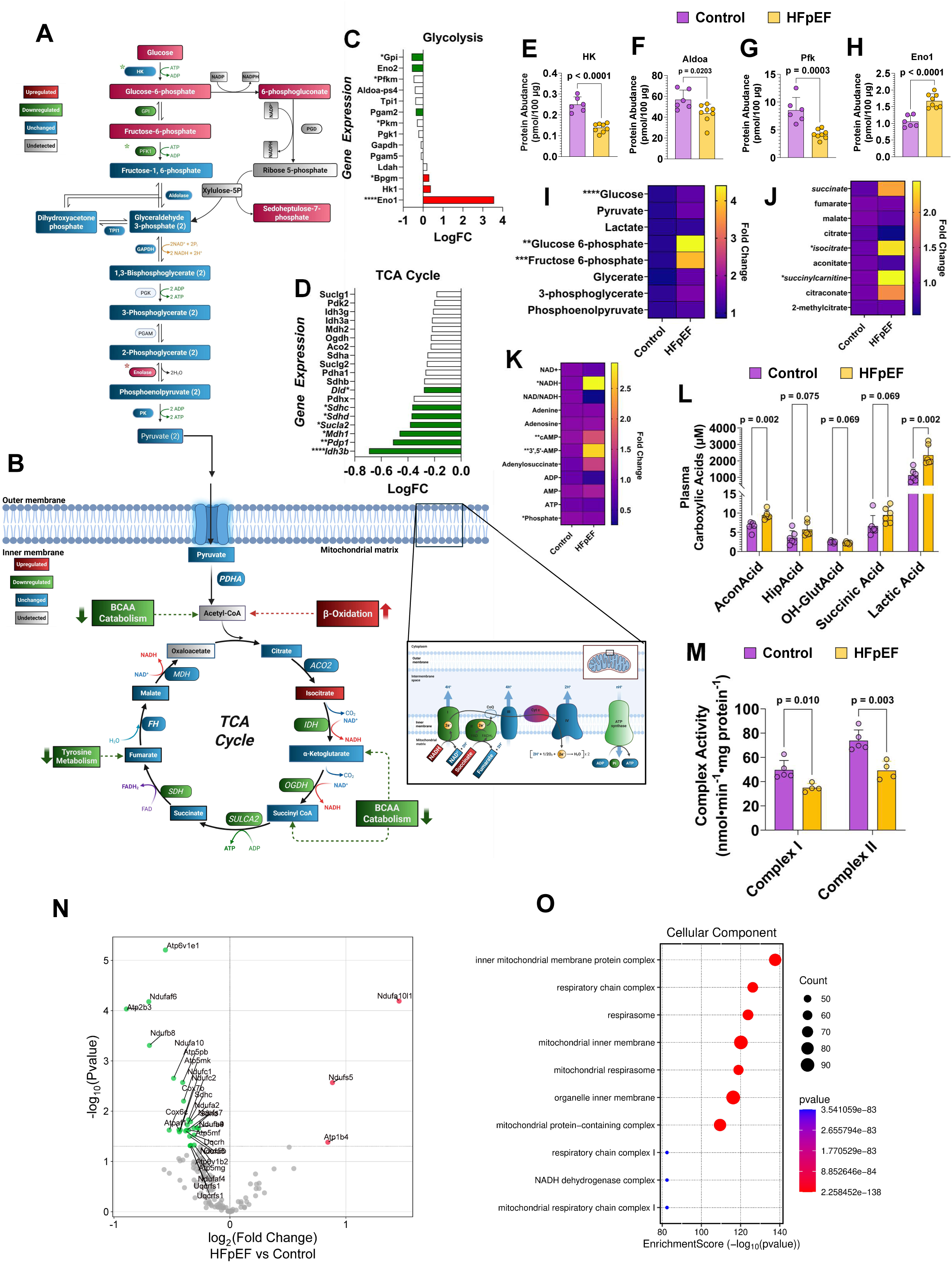
Reduced Glycolysis and TCA Cycle Activity in HFpEF Skeletal Muscle. Integration of metabolites, gene/proteins involved in **(A)** glycolysis and **(B)** TCA cycle. Astrix indicates that for a given gene, the corresponding protein is also altered according to the previously mentioned color scheme. Log fold changes for differentially regulated genes for **(C)** glycolysis and **(D)** TCA cycle. **(E-H)** Quantification of proteomics for glycolysis-related proteins. Metabolites related to **(I)** glycolysis, **(J)** TCA cycle, and **(K)** cofactor metabolism were altered between HFpEF and control skeletal muscle. **(L)** Plasma carboxylic acids. **(M)** Skeletal muscle (tibialis anterior) Complex I and II activity. **(N)** Volcano plot **(O)**, dotplot of mitochondrial genes curated from the mitocarta database (version 3.0). All data are mean ± SD. Adjusted p-values are reported for significantly altered genes **(C & D)**. An unpaired t-test was used to determine the difference between (**E-H, L-M**). Metabolites **(I-K)** were analyzed with a Welch’s two-sample t-test. n=6-8/group. Significant values are provided within graphs or for gene expression and metabolite accumulation *P<0.05, **P<0.01, ***P<0.001, ****P<0.0001. Red and green indicate up- and down-regulation of genes and metabolites, respectively. Blue and silver indicate unchanged and undetected genes and metabolites, respectively.

### Skeletal Muscle BCAA Catabolic Defects

A BCAA catabolic defect is noted in cardiac metabolism in patients with HFpEF and HFrEF, as well as the ZSF1 obese rat^24,49^. Whether skeletal muscle BCAA metabolism is also impacted by HFpEF is not fully known. Gene expression and proteins associated with BCAA catabolism were mostly downregulated in HFpEF (Figure 3A-C). Despite no observed difference in the phosphorylation state of branched-chain keto acid dehydrogenase kinase subunit alpha (BCKDHA) (Supplemental Figure 4). BCAAs (leucine, isoleucine, and valine) were increased in the gastrocnemius (Figure 3D), soleus, and EDL skeletal muscles (Supplemental Figure 5). Targeted analysis of the plasma confirms a systemic accumulation of BCAAs (isoleucine and valine) (Figure 3E) in HFpEF. Finally, we targeted the ratio of C5-OH carnitine and C3-DC (dicarboxylic) because it is considered a reliable biomarker of tissue BCAA catabolism^50^. Figure 3F shows a significant reduction of BCAA-catabolic specific metabolites C5-OH/C3-DC ratio in plasma, soleus, and EDL muscle in HFpEF rats. Herein, we provide an unambiguous metabolic signature of defective BCAA catabolism in HFpEF skeletal muscle.

**Figure 3.**
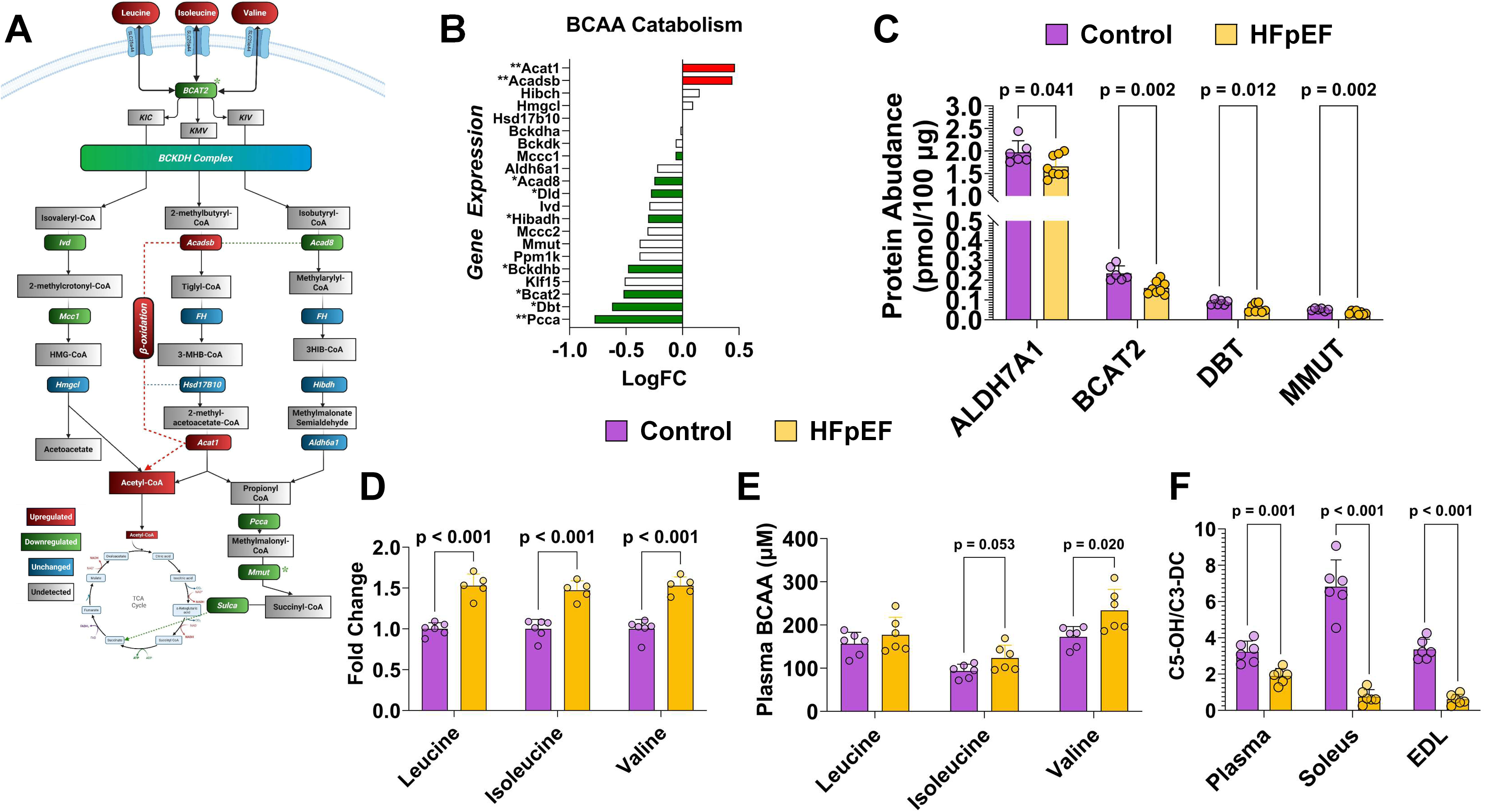
Branched Chain Amino Acid (BCAA) Catabolic Defect in HFpEF Skeletal Muscle. **(A)** Visual integration of BCAA metabolism. **(B)** Differential gene expression (C) quantified proteomics of proteins in BCAA metabolism. **(D)** Skeletal muscle and plasma **(E)** BCAA amino acid. (F) The ratio of C5-OH and C3-DC acylcarnitines from targeted metabolomics. All data are mean ± SD. An unpaired *t*-test was used to determine significance for figures **B-H**. Adjusted p-values are reported for significantly altered genes **(B)**. N=6-8/group. Significant values are provided within graphs or for gene expression *P<0.05, **P<0.01, ***P<0.001, ****P<0.0001. Red and green indicate up- and down-regulation of genes and metabolites, respectively. Blue and silver indicate unchanged and undetected genes and metabolites, respectively. Astrix indicates that for a given gene, the corresponding protein is also altered according to the previously mentioned color scheme.

### Altered Tryptophan-Kynurenine-NAD^+^ Axis in HFpEF Skeletal Muscle

Tryptophan metabolism was also highly represented in our pathway-level analysis. The relationship between tryptophan and its metabolites, kynurenine and kynurenic acid (i.e., kynurenate), has been associated with physical function and activity, making this pathway particularly relevant for exercise intolerance in HFpEF^48^. Gene expression associated with tryptophan breakdown was mostly upregulated (Figure 4A) in HFpEF skeletal muscle. The 2.3-fold increase (adjusted p=0.01) in Tdo2 gene expression is particularly relevant as this encodes the rate-limiting enzyme in tryptophan metabolism, which converts tryptophan to kynurenine. Kyat3 is an amino transferase that functions to convert kynurenine to functional kynurenic acid. Gene expression shows a reduction in Kyat3 and Kyat 1, which was supported by proteomics analysis (Figure 4B), also demonstrating a reduction in KYAT1. The differential gene expression of Tdo2 and Kyat3 suggests that the kynurenine-serotonin axis is altered in HFpEF skeletal muscle. Despite the increase in Tdo2, our metabolomics analysis (Figure 4C) demonstrates reduced levels of kynurenine (-0.88 log2FC, p=0.006) and kynurenate (-0.67 log2FC, p=0.01), whereas serotonin was increased (0.53 log2FC, p=0.01) in gastrocnemius muscle. We used the ratio of kynurenine to tryptophan as a proxy to determine the activity of rate rate-limiting enzyme of tryptophan metabolism, idoleamine-2,3-dioxygenase (IDO), as previously reported^51^. IDO activity was reduced in gastrocnemius and EDL tissues with no change in plasma or soleus tissue (Figure 4D). Similar to the increase in serotonin in the gastrocnemius (Figure 4E), we also show increased serotonin in the EDL muscle. Contrary to the EDL, the soleus muscle showed reduced serotonin, indicating differential dysregulation in tryptophan metabolism between glycolytic and oxidative muscle groups. Plasma serotonin was not different between control and HFpEF (3.14 ± 3.5 vs. 3.92 ± 3.9 µM), suggesting that levels of skeletal muscle serotonin are due to endogenous synthesis rather than circulating serotonin. We expanded our metabolomics analysis to the plasma to target tryptophan-specific metabolites. Figure 4F demonstrates that plasma tryptophan is reduced along with indole-3-propionic acid (3-IPA) and indoxyl sulfate (Ind-SO4), but increased plasma indole-3-acetic acid (3-IAA). Other metabolites (indolelactate and indolproprionate) associated with microbial metabolism of tryptophan were also accumulated in HFpEF skeletal muscle (0.48 Log2FC, p<0.0001 & 0.48 Log2F, p=0.01).

**Figure 4.**
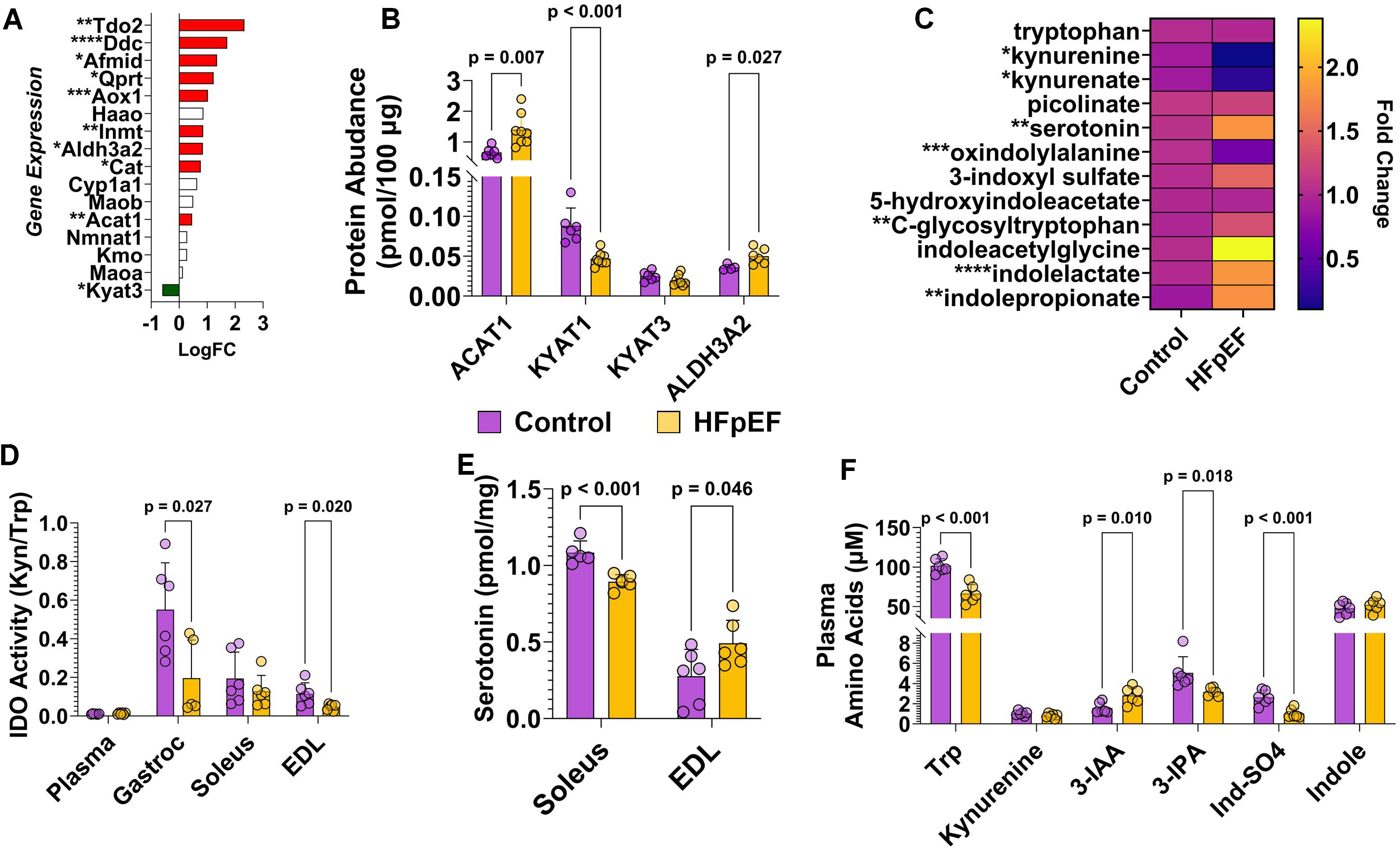
Tryptophan metabolism favors serotonin formation in HFpEF skeletal muscle. **(A)** Tryptophan metabolism gene expression (LogFC) and **(B)** quantification of proteomics. **(C)** Heatmap of tryptophan-related metabolites. **(D)** Ratio of kynurenine versus tryptophan as an index of IDO enzyme activity. **(E)** Skeletal muscle (soleus and EDL) serotonin concentration from targeted metabolomics. **(F)** Plasma tryptophan and related metabolites. All data are mean ± SD. An unpaired *t*-test was used to determine significance for figures B-H Adjusted p-values are reported for significantly altered genes. N=6-8/group. Significant values are provided within graphs or for gene expression and metabolite accumulation *P<0.05, **P<0.01, ***P<0.001, ****P<0.0001.

### Inefficient β-Oxidation Defines the Metabolic Myopathy Signature in HFpEF

Fatty acid metabolism is the predominant substrate in skeletal muscle at rest and during exercise. Clinical evidence suggests that the capacity to oxidize fatty acids is impaired in HFpEF skeletal muscle^11^. We demonstrate a broad increase in genes, proteins, and metabolites associated with β-oxidation in HFpEF skeletal muscle (Figure 5A). At the gene and protein level (Figure 5B), acyl-CoA synthetase enzymes (ACSL3 and ACSL6) were reduced in HFpEF skeletal muscle. Only the expression of Acsl4 was increased significantly in HFpEF. Proteins involved in acylcarnitine/acetylcarntine synthesis and transport (Figure 5C) (i.e., CrAT and SLC25A20) were upregulated. Acyl-CoA synthetase enzymes (ACADL, ACADS, and ACADSB) and hydratase (ECH1, ECI1, and ECI2) were increased in HFpEF skeletal muscle. Enoyl-CoA hydratase domain-containing 2 (Echdc2) was uniquely decreased at both the gene and protein levels. Echdc2 is not a canonical β-oxidation enzyme, but is structurally similar to Echdc1, which maintains fidelity of the acyl-CoA pool by repairing acyl-CoA metabolites such as methylmalonyl-CoA^52^. Some evidence suggests Echdc2 regulates LCFA biosynthesis, while other work highlights its role in the degradation of 3-methylcrotonyl-CoA carboxylase (Mccc2) to control BCAA metabolism^53^. Echdc2 is potentially a critical signaling node that, when reduced, diminishes the fine-tuning of β-oxidation and BCAA metabolism to adequately metabolize unique acyl-CoA species.

**Figure 5.**
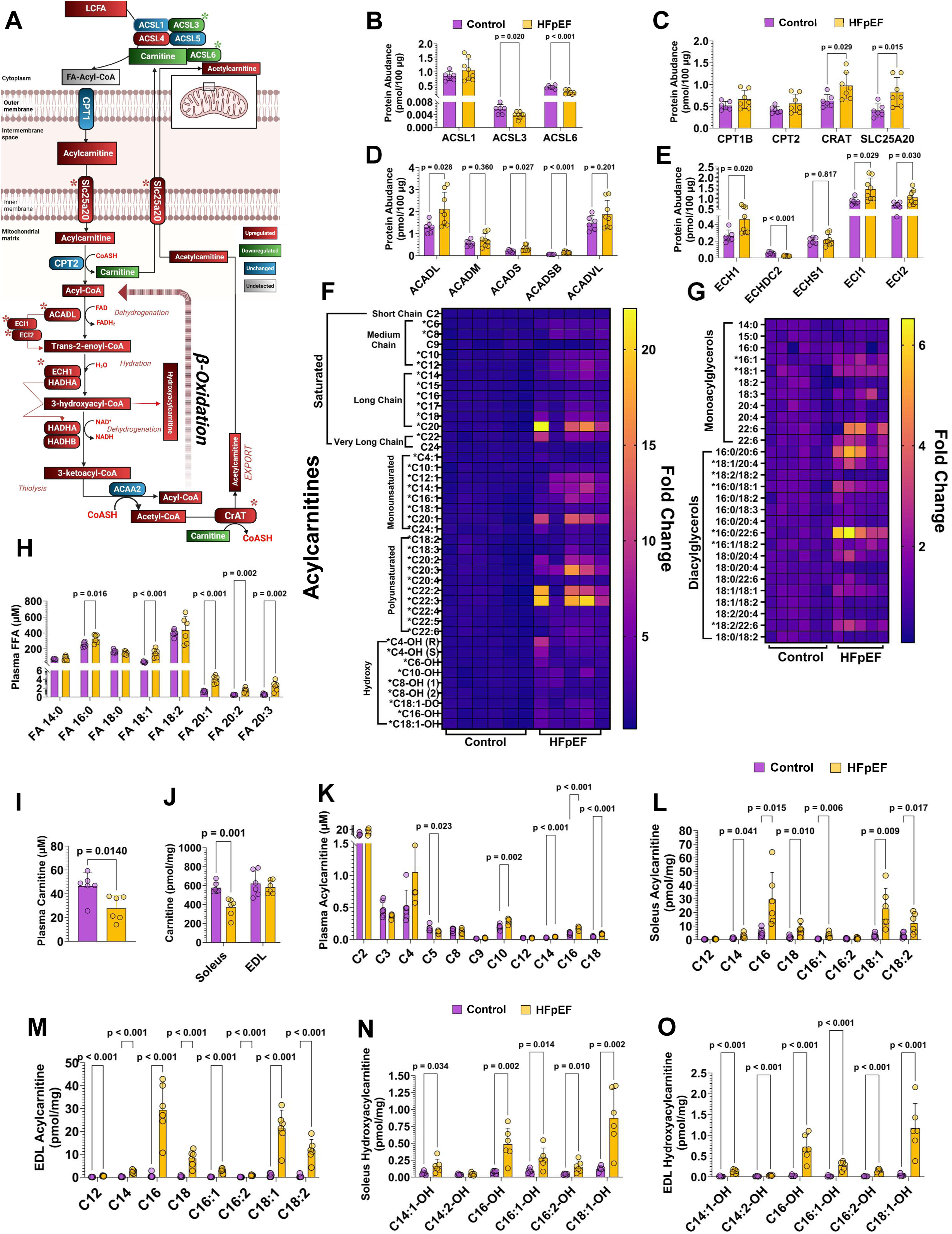
Upregulated but Incomplete β-oxidation Drives Acylcarnitine Synthesis in HFpEF Skeletal Muscle. **(A)** Visual integration of genes and metabolites for β-oxidaiton. Astrix indicates that for a given gene, the corresponding protein is also altered according to the previously mentioned color scheme. Quantification of proteomics analysis for β-oxidaiton proteins involved in **(B)** activation of LCFA and the conversion to fatty acyl-CoA, **(C)** fatty acid transport, **(D)** acyl-CoA dehydrogenation, and **(E)** peroxisomal and mitochondrial double-bond processing. Heatmap of skeletal muscle **(F)** acylcarnitine and **(G)** diacyl-and monoacylglycerol species from global metabolomic profiling. **(H)** Plasma free fatty acids. **(I)** Plasma and **(J)** skeletal muscle free carnitine levels. Quantified **(K)** plasma, **(L)** soleus, and **(M)** EDL long-chain acylcarnitines. **(N)** Soleus and **(O)** (EDL) hydroxyacylcarnitine concentrations. All data are mean ± SD. An unpaired *t*-test was used to determine significance for figures B-H & I-O. Adjusted p-values are reported for significantly altered genes. N=6-8/group. Significant values are provided within graphs. Red and green indicate up- and down-regulation of genes and metabolites, respectively. Blue and silver indicate unchanged and undetected genes and metabolites, respectively.

Figure 5F demonstrates a significant accumulation of saturated and unsaturated acylcarnitine species in the skeletal muscle HFpEF rats. Long chain (C14-C20) and very long chain (>C20) showed the highest levels of accumulation (Supplemental Table 2.1). Similarly, we show accumulation of mono- and diacylglycerols with a propensity for accumulation of long-chain and very-long-chain species (Figure 5G). The substantial accumulation of acylcarnitines and metabolites is associated with incomplete β-oxidation, suggesting a systemic overload of circulating fatty acids. Plasma metabolomics confirms increased circulating long-chain fatty acids (LCFA) (16:0, 18:1, 20:1, 20:2, and 20:3) (Figure 5H). We also confirm that global carnitine bioavailability (required for activation of fatty acids) is reduced (Figure 5I). Carnitine levels in the soleus, but not the EDL, were depleted in HFpEF (Figure 5J). We followed up on our global metabolomic and lipidomics analysis to target specific acylcarnitines in plasma, soleus, and EDL muscles. Saturated (Figure 5K) and unsaturated (Supplemental Figure 6) long-chain acylcarnitines were increased in the plasma. Likewise, both the soleus and EDL showed robust accumulation of C16, C16:1, C18, and C18:2 long-chain acylcarnitines (Figures 5L-M). Our pathway analysis shows that β-oxidation generates increased levels of 3-hydroxyacyl-CoA and the generation of highly toxic hydroxyacylcarnitines. Targeted lipidomics confirms elevated long-chain hydroxyacylcarntines accumulation in both oxidative soleus (Figure 5O) and glycolytic EDL (Figure 5P) skeletal muscle. Collectively, these data demonstrate several points of dysregulation in β-oxidation in HFpEF skeletal muscle.

### Wide-spread Phospholipid Remodeling is a Key Feature in Cardiometabolic HFpEF

Phospholipids constitute the majority of the cell membranes and are essential regulators of signal transduction, insulin sensitivity, and force production in skeletal muscle. PPAR signaling is an important upstream regulator of phospholipid metabolism because it is involved in the regulation of lipid transport and adipocyte differentiation. PPAR signaling genes (CD36 & FABP) were highly upregulated in the gastrocnemius of the HFpEF rats (Figure 6A-B). Gene expression analysis of the glycerophospholipid metabolism pathway shows a robust upregulation of phospholipase Plag2g7 and a downregulation of Gpcpd1 in gastrocnemius skeletal muscle (Figure 6C). Plag2g7 is one of 61 genes that are commonly upregulated (Adjusted P<0.05) between gastrocnemius, soleus, and EDL skeletal muscle in HFpEF skeletal muscle (Supplemental Table 1.8). Proteomics confirms this reduction in GCPD1 protein as well as the choline transporter SLC44A2 (Figure 6D). Choline is an essential nutrient and a direct precursor for the membrane lipids phosphatidylcholine and sphingomyelin^54^. Interestingly, choline was significantly increased, despite the decrease in Slc44a2, along with several choline-associated phospholipids (Figure 6E). Among these choline metabolites was trimethylamine-N-oxide (TMAO), which is a toxic metabolite generated in the liver from the oxidation of gut-born TMA^55^. While skeletal muscle cannot directly synthesize TMAO, it is capable of uptake from the circulation. Our targeted analysis confirms elevated plasma choline and TMAO, while an important inhibitor of TMAO, trigonelline, is reduced in HFpEF (Figure 6F). Figures 6G-H show that choline is only increased in the EDL, whereas TMAO is only significantly increased in the soleus muscle. Trigonelline is reduced in both muscles in HFpEF. Figure 6I provides a comprehensive heatmap of phosphoipids from gastroneiums samples. Lysophosphatidylcholine (LPC) is formed from phosphatidylcholine and is a key component of oxidized low-density lipoproteins^56^. Our analysis reveals a significant accumulation of lysophosphatidylcholine and lysophosphoethanolamine. Whereas some LPCs such as (1-nonadecanoyl-GPC (19:0), 1-arachidoyl-GPC (20:0), 1-eicosenoyl-GPC (20:1) 1-docosahexaenoyl-GPC (22:6), 1-lignoceroyl-GPC (24:0) were significantly reduced. Of the 46 phosphatidylcholine species detected, 18 species were reduced, with 10 species increased (Supplemental Table 2). Sixteen of the 23 phosphoenathalonamines (PE) were increased in the skeletal muscle, with no species being reduced. Phosphatidylglycerols (PG) species and phosphatidylinositol (PI) were mostly increased. Whereas phosphatidylserine (PS) species were mostly reduced except for 1-stearoyl-2-dihomo-linolenoyl-PS (18:0/20:3n3 or 6). Contrary to the accumulation of most lipid species in HFpEF skeletal muscle, we demonstrate that plasmalogens, a glycerolphospholipid generated within peroxisomes and the endoplasmic reticulum, are dramatically reduced in HFpEF skeletal muscle (Figure 6J).

**Figure 6.**
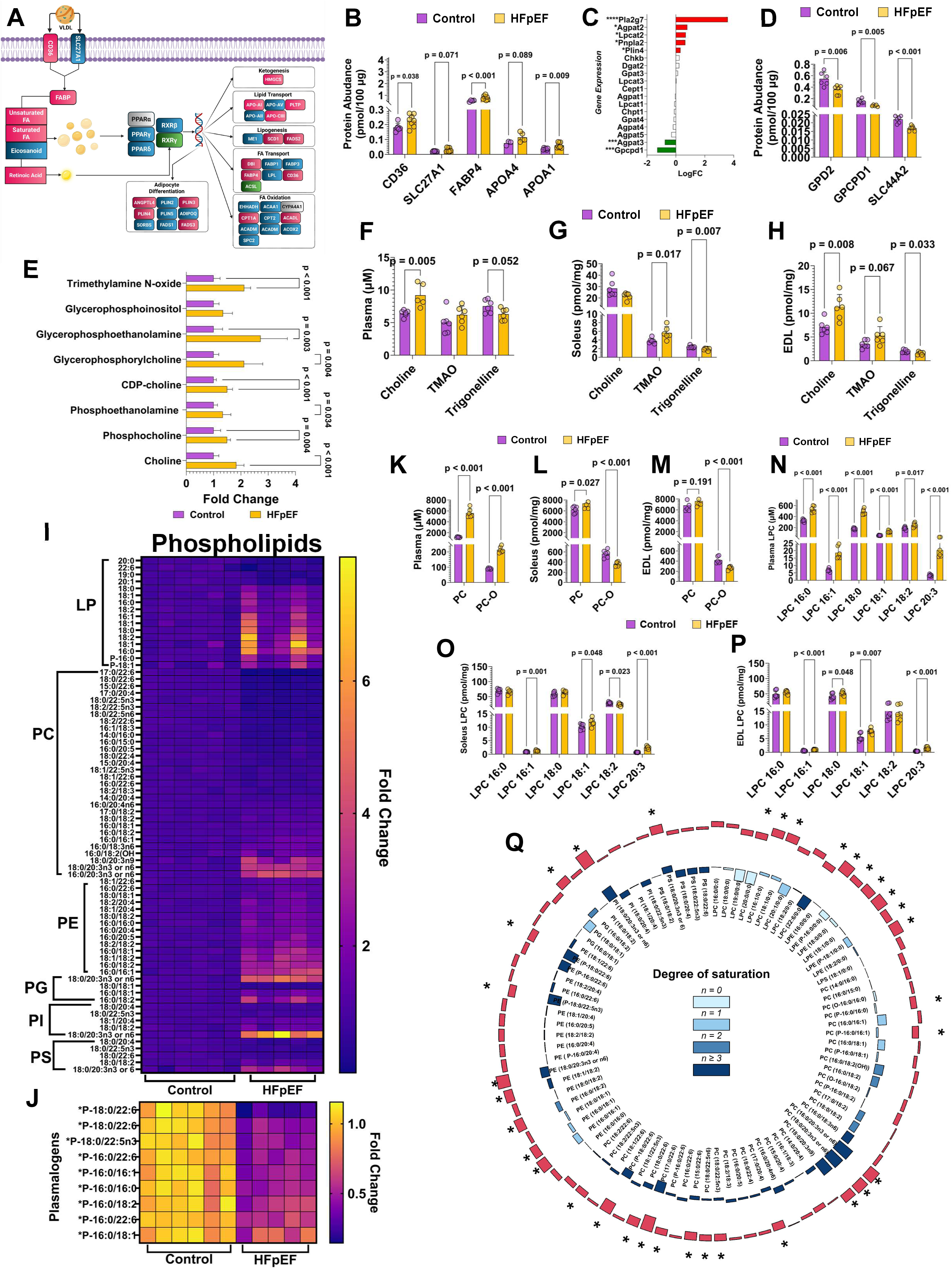
Pla2g7 remodels phospholipid metabolism in HFpEF skeletal muscle. **(A)** Visual integration of PPAR signaling. **(B)** Fatty acid transport proteins. **(C)** Gene expression and **(D)** protein quantification of glycerophospholipid metabolism. **(E)** Skeletal muscle (gastrocnemius) metabolites related to glycerophospholipid and choline metabolism. (F) Plasma, (G) soleus, and (EDL) choline TMAO, and trigonelline concentrations. (H) Heatmap of global lipidomics for all phospholipids and (J) plasmalogens. (K) Plasma, (L) soleus, and (M) EDL phosphatidylcholine (PC) and PC-O species. (N) Plasma, (O) soleus, and (P) EDL lysophosphatidylcholine (LPC) concentrations. (Q) Circos plot for degrees of saturation/unsaturation of phospholipid species from global lipidomics. Red bars represent the total fold change, and blue shades represent the degree of saturation. All data are mean ± SD. An unpaired *t*-test was used to determine significance for figures B, D, F, G, H, K-P. Adjusted p-values are reported for significantly altered genes. n=6-8/group. Significant values are provided within graphs or for gene expression *P<0.05, **P<0.01, ***P<0.001, ****P<0.0001. Lyosphospholipids (LP), phosphatidylcholine (PC), phosphatidylethanolamines (PE), phosphoglycerol (PG), phosphoinositols (PI), and phosphotidylserine (PS).

We next sought to determine if the PC abundance was reflected in the circulation and if skeletal muscle with different fiber type profiles was impacted in HFpEF. We targeted the sum of 38 standard PC species and 40 PC vinyl-ether-linked PC species to evaluate lipids generated by the peroxisome. Plasma PC and alkyl-ether (O) PC-O were both increased in the plasma (Figure 6K). The soleus showed an increase in PC (Figure 6L), whereas there was no difference between control and HFpEF in the EDL (Figure 6M). The PC-O content was significantly reduced in both the soleus and EDL, suggesting a reduction in PC generated by the peroxisome. Similarly, we quantified the LPC species. Increases in plasma (Figure 6N) were mirrored in the soleus (Figure 6O) and EDL muscle (Figure 6P). Shingomyelin and ceramides are structurally related but distinct sphingolipid species. We demonstrate that proapoptotic ceramides were mostly increased in plasma and skeletal muscle (Supplemental Figure 7 & Table 2.3). Whereas, shingomyelin, an important structural sphingolipid, is elevated in the plasma but significantly reduced in soleus and EDL muscles (Supplemental Figure 8 & Table 2.4). Figure 6Q summarizes the shifting composition of phospholipid species in skeletal muscle, favoring increased saturation of phospholipids and increasing arachonyl lipid species.

## Discussion

HFpEF is a debilitating chronic form of heart failure that presents with multiple phenotypes. The obesity epidemic has converged with HFpEF to produce a severe cardiometabolic phenotype marked by profound exercise intolerance. Exercise intolerance is attributed to both cardiac and non-cardiac origins in HFpEF. Recent reports have described transcriptional, proteomic, and metabolomic signatures of cardiac dysfunction in cardiometabolic HFpEF^24,57,58^. However, the pathways that mediate skeletal muscle dysfunction in HFpEF are not well understood. Therefore, the field lacks the critical insight into this key target for therapeutic intervention. Our goal was to elucidate the molecular signature of skeletal muscle pathobiology in cardiometabolic HFpEF. We build on previous observations of exercise intolerance (Figure 1A), muscle weakness (Figure 1B), fiber atrophy, interstitial fibrosis, and reduced capillary density (Figure 1F-I). We also provide new evidence of adipose infiltration (i.e., myosteotosis) into the skeletal muscle (Figure 1J). This is an important distinction from other studies that have shown increased intracellular lipid accumulation in this model^59^. Intramyocellular lipids refer to the storage of neutral ectopic lipids that are associated with insulin resistance, but have also been shown to be highly enriched after long-term endurance exercise training^60^. Therefore, the presence of intramyocellular lipids does not indicate metabolic dysfunction and poor muscle performance. Whereas intramuscular adipose tissue accumulation, as we demonstrate, is directly related to metabolic dysfunction, muscle weakness, and impaired exercise capacity in HFpEF^15,61^. This finding is highly significant because it confirms the ZSF1 obese rat is indeed a translational model of skeletal muscle dysregulation in HFpEF.

At the transcript and protein levels, we show that major energy-producing pathways (TCA cycle, OXPHOS, ETC) are downregulated in HFpEF skeletal muscle. Consistent with the insulin resistance phenotype of the ZSF1 obese rat, we show impaired glucose metabolism (Figure 2A). The downregulation of glycolysis occurs at the level of the rate-limiting enzyme phosphofructokinase (Pfk). The enzyme α-enolase (Eno1) was unique because it was the only glycolytic enzyme highly upregulated in the skeletal muscle (Figure 2C & H). However, its participation in the conversion of 2-phosphoenolglycerate to phosphoenolpyruvate is likely mitigated by its alternative function to act as a cell-surface receptor for plasminogens and the deleterious remodeling of the cytoskeleton^62^. The reduction in TCA cycle gene expression (Figure 2D) with concordant accumulation of isocitrate (Figure 2J) and reducing equivalents (Figure 2K) suggests an inability of the TCA cycle to support ATP synthesis. The accumulation of TCA intermediates in the plasma supports this systemic impact of skeletal muscle metabolism in HFpEF. While mitochondrial dysfunction has generally been described in HFpEF, there is little specific knowledge of the key nodes involved in skeletal muscle. Our data reveal that reduced complex I and II activities are key sites that potentially explain the accumulation of NADH and succinate in the skeletal muscle (Figure 2B). Complex I dysfunction appears to be a major contributor to increased reductive pressure in skeletal muscle, with approximately 40% of complex I assembly genes reduced in HFpEF muscle. Our data adds biological insight into the nature of mitochondrial dysfunction in HFpEF, supporting previous work showing reduced citrate synthase activity and mitochondrial respiration^21,63^.

In both HFpEF and HFrEF, BCAA metabolism has garnered much attention because of its prominent role in insulin signaling and mitochondrial function. The BCAAs (leucine, isoleucine, and valine) are essential amino acids whose carbon skeletons can be propelled into the TCA cycle under energetically demanding conditions^64^. Previous reports demonstrate that improving BCAA catabolism can improve cardiac function in HFrEF^65^. Recent work has demonstrated that in patients with HFpEF, myocardial BCAA levels are increased and BCAA catabolic products are decreased when compared to healthy individuals and patients with HFrEF^24^. Collectively, these reports indicate the myocaridal BCAA catabolic defect is a defining molecular feature in HFpEF.

While cardiac BCAA metabolism is regarded as an important metabolic signature of heart failure, skeletal muscle is the predominant site of *in vivo* BCAA catabolism^66^. During exercise, BCAAs are diverted towards oxidation and away from protein synthesis^64^. The ablation of branched-chain aminotransferase (Bcat2) in the skeletal muscle reduced exercise performance and shifted substrate utilization in mice^67^. We demonstrate that, like myocardial tissue, skeletal muscle also shows a similar BCAA catabolic defect (Figure 3A). While the global signatures are similar, we show that Bcat2 gene expression and protein abundance are reduced in the skeletal muscle, whereas this transaminase is increased in HFpEF myocaridum^24^. Because the BCAAs are obtained through the diet, and ZSF1 obese rats are hyperphagic, we targeted a reliable metabolite ratio (C5-OH/C3-DC) consistent with BCAA oxidation^50,68,69^. Our results differ from previous reports, which show increased BCAA catabolism in the skeletal muscle of Zucker Diabetic Fatty (ZDF) rats^50^. Our use of the ZSF1 rat, which is derived from the crossing of a ZDF female rat with a male spontaneously Hypertensive Heart Failure (SHHF) rat, sheds mechanistic insight into distinguishing metabolic dysregulation associated with obesity without HFpEF (ZDF rat) versus cardiometabolic HFpEF (ZSF1 obese rat). The impact of hypertension alone without obesity, when comparing ZSF1 lean rat skeletal muscle to WKY control, does not alter BCAA gene expression or protein abundance (Supplemental Table 1.2). Rather, the synergistic combination of obesity and hypertension appears to promote the BCAA catabolic defect in ZSF1 obese rats regardless of the control strain (Supplemental Table 1.3). Given the dominant role of skeletal muscle in global BCAA oxidation, the abundance of BCAA measured in the plasma of HFpEF patients likely reflects skeletal muscle rather than myocardial BCAA catabolic defects.

Recently, plasma tryptophan has emerged as a potential biomarker for low physical capacity in older individuals. In older (68.3 ± 1.0 years) participants who maintain high levels of physical activity and regular endurance exercise, skeletal muscle abundance of tryptophan metabolites, kynurenine, and kynurenate was increased when compared to age-matched sedentary individuals^51^. These metabolites were highly associated with cardiorespiratory fitness and mitochondrial function. Kynurenate bioavailability promotes quinolinic acid synthesis and the formation of NAD^+70,71^. Our findings demonstrate a novel phenomenon in cardiometabolic HFpEF whereby skeletal muscle tryptophan metabolism is diverted away from kynurenine, kynurenate, and subsequent NAD^+^ formation towards serotonin synthesis. The loss of the tryptophan-kynurenine-NAD^+^ axis could potentially mediate the low NAD^+^/NADH ratio we observed (Figure 2K). In this case, the diversion towards serotonin synthesis likely contributes further metabolic dysregulation in the skeletal muscle. Previous studies demonstrated promising effects in blocking serotonin signaling to reduce metabolic dysfunction associated with obesity^72^. The accumulation of skeletal muscle serotonin also has implications for skeletal muscle quality and function, as it is involved in myogenic differentiation and excitation-contraction^73^. The receptor for serotonin (5-HT_2A_) is localized to the t-tubules, where it can trigger calcium release from the sarcoplasmic reticulum to induce muscle contraction by increasing the release of acetylcholine^73^. Some evidence suggests that local skeletal muscle conversion of tryptophan to serotonin is increased during intense exercise and is required for adaptation to exercise training^74^. However, in this context, ZSF1 obese rats have severe exercise intolerance and do not engage in high levels of physical activity. Our results align with earlier studies implicating serotonin in skeletal muscle myopathy related to its postsynaptic action to alter membrane polarization, resulting in muscle force production. More mechanistic studies are required to fully elucidate whether intramuscular serotonin is involved in skeletal muscle weakness in HFpEF.

Fatty acids are the primary source of ATP during sustained low-to-moderate intensity exercise. In HFpEF patients, skeletal muscle oxidative capacity is substantially impaired^11^. The degree of metabolic dysfunction is closely associated with reduced exercise capacity^75^. Despite these reports, there is little insight into the major signaling nodes and metabolic fingerprint of HFpEF skeletal muscle. Acylcarnitines function to transport fatty acid acyl groups from the cytosol to the mitochondria to undergo β-oxidation. The accumulation of acylcarnitines in the plasma is an established biomarker for obesity and diabetes. Hunter et al (2016) reported that in HFpEF patients with obesity, plasma long-chain acylcarnitines are increased^76^. Follow-up studies using a larger cohort with greater levels of obesity (39.8 kg/m^2^) showed no increase in plasma acylcarnitines in HFpEF patients^24^. Myocardial metabolomics in the same study showed a marked reduction in acylcarnitines in HFpEF versus HFrEF patients. Our reports of elevated acylcarnitines in skeletal muscle of ZSF1 obese rats align more closely with a recent report from Gibb et al (2025) that who reported a similar increase in acylcarnitines from the left ventricle of ZSF1 obese rats ^77^. While acylcarnitines are elevated in both tissues, our work shows that the transcription and protein regulation of fatty acid oxidation are increased in skeletal muscle; this same program was downregulated in myocardial tissue. The discrepancy between reports of increased versus decreased acylcarnitine in HFpEF is potentially related to the progression and severity of cardiometabolic HFpEF. As the degree of obesity and HFpEF develops over time, the capacity to generate acylcarnitines is presumably diminished, as is fatty acid oxidation. It is intriguing to speculate if skeletal muscle follows a parallel trajectory with myocardial metabolism.

The skeletal muscle is one of the predominant sources of circulating acylcarnitines, and based on its mass, plasma acylcarnitines are most likely a reflection of inefficiency in skeletal muscle fatty acid oxidation. Many potential explanations possibly explain the discrepancy between plasma and tissue acylcarnitine in HFpEF. Carnitine (L-carnitine) is a critical metabolite required for the formation of acylcarnitines via the actions of CPT1b to free carnitine to long-chain fatty acids for transport across the mitochondrial membranes^78^. Carnitine buffers excess acyl-CoA species and maintains the free CoA pool. Subsequent analysis from other groups involving a larger number of HFpEF patients demonstrates significant reductions in plasma carnitine bioavailability when compared to non-failing controls and HFrEF patients^79^. We confirm in the ZSF1 obese model a 45% reduction in plasma carnitine bioavailability (Figure 5I). The highly oxidative soleus muscle was similarly depleted of carnitine (Figure 5J). While our results show clear reductions in carnitine bioavailability, the mechanistic link between carnitine and dysregulation of fatty acid oxidation is unclear. The abundance of long-chain acylcarntine species provides evidence that carnitine is being utilized for activation, but β-oxidation is not capable of complete breakdown. The actions of carnitine acetyltransferase (CrAT) further deplete the supply of carnitine by generating acetylcarnitine for export to relieve lipid stress. In the current circumstance, the influx of long-chain fatty acids depletes free carnitine to buffer acyl-CoA at the expense of the free CoA pool. This is an important consideration because there is some evidence linking cardiac dysfunction in heart failure to reduced CoA levels^79^. Our current findings suggest that CoA may also be depleted in skeletal muscle and may contribute to the dysfunction in HFpEF. Future work is required to determine if CoA dysregulation is a causative factor in skeletal muscle dysfunction in HFpEF.

Global lipidomics (Figure 5F) identified several hydroxyacylcarntine (Acyl-OH) species that increase in HFpEF skeletal muscle. We verified in both the soleus and EDL muscles numerous hydroxyacylcarnities (Figure 5N-O). These toxic acylcarnitine species are in response to increased production β-hydoroxylacyl-CoAs via β-hydoroxylacyl-CoA dehydrogenase and likely reverse efflux through CPT2. The position of the OH-group attached to the β-carbon can promote an altered NADH: NAD^+^ ratio and reduce β-oxidation efficiency, which is consistent with our findings (Figure 2K)^78^. The elevation of this rare acylcarnitine species is associated with mitochondrial myopathies and is positively associated with the mutational load of disease severity^80^. Other reports in mitochondrial DNA mutations also implicate hydroxyacylcarnitines as a driver of increased reductive stress^53^. Our current report provides new evidence of a similar myopathy signature and further defines the state of mitochondrial dysfunction in cardiometabolic HFpEF.

Fatty acid overload is mechanistically implicated in skeletal muscle dysfunction in HFpEF. Using a “two-hit” (high-fat diet and L-NAME) murine model, we demonstrated transcriptional upregulation of fatty acid transport genes in the skeletal muscle when compared to control mice^81^. We demonstrate a consistent signature in the robust ZSF1 obese model of cardiometabolic HFpEF. The activation of PPAR signaling promotes the influx of lipid transport and lipogenesis genes. Ether lipid and glycerolphospholipid metabolism were highly enriched pathways (KEGG, NES=1.7, p=0.003) related to lipid synthesis. Our novel findings demonstrate that the activation of phospholipases (Pla2g7) breaks down membrane phospholipids to generate LPC (Figure 6N-M) and oxidized fatty acids. The upregulation of Pla2g7 also increases the breakdown of (arachidonic acid) AA- and (docosahexaenoic acid) DHA-containing PCs, which alters the saturation state of the outer phospholipid membrane. We also show a strong signaling for increased free choline accumulation in the skeletal muscle. The downregulation of the phosphodiesterase Gpcpd1 reduces glycerolphosphocholine (GPC) hydrolysis, therefore increasing GPC accumulation (Figure 6E). The impact of increased Pla2g7 and decreased Gpcpd1, combined with elevated choline levels, suggests that choline cycling is disrupted in HFpEF skeletal muscle. Excessive choline can promote inflammation through the activation of toll-like receptors via LPC and the conversion to TMAO. Plasma choline levels were increased in ZSF1 obese rats, which may drive increased TMAO uptake and contribute to the inflammatory profile of skeletal muscle in HFpEF. The lipid composition of the phospholipid membrane tracks with our observation of increased expression of ACSL4, which prefers ω-6 fatty acids (AA., 20:4) and reduced expression of ACSL6, which is selective for ω-3 fatty acids (DHA, 22:6). Our lipidomics analysis shows this altered signaling cascade remodels the phospholipid membrane by increasing AA and decreasing DHA incorporation into PC and PE. The increased ω-6: ω-3 ratio skews the phospholipid membrane towards a pro-inflammatory state, making the membrane more prone to lipid peroxidation (i.e., ferroptosis). The collective impact of this phospholipid remodeling requires additional focus. Our results strongly suggest that phospholipid membrane fluidity, signaling transduction, and stability are impaired in HFpEF skeletal muscle. The liberation of LPC from PC-membrane via increased Plag2g7 is a novel mechanism linking skeletal muscle phospholipid metabolism to systemic inflammation in the setting of cardiometabolic HFpEF. The role of Pla2g7 in skeletal muscle has yet to be elucidated. Systemic deletion of Pla2g7 protects aged mice from metabolic dysfunction and chronic inflammation when fed a high-fat diet, suggesting a deleterious role in obesity and aging^82^. Interestingly, analysis of high and low-functioning octogenarians suggests a positive association between serum LPC and physical fitness^48^. Our results strongly oppose this finding, which could be related to the highly obese condition of our model. Future work is required to determine if Pla2g7 and LPCs are consistent biomarkers of skeletal muscle dysfunction and exercise intolerance in cardiometabolic HFpEF.

Our data clearly demonstrates both mitochondrial and peroxisomal overload in HFpEF skeletal muscle. Peroxisomes generate medium, branched-chain, and many of the rare acylcarnitine species (dicarboxyl, DC) were detected in our analysis. The peroxisome performs its own β-oxidation to generate shorter-chain fatty acids for the mitochondria for β-oxidation^83^. Peroxisomal β-oxidation provides shorter acetyl groups specifically used to synthesize malonyl-CoA, indicating an important role for these metabolites in negatively regulated fatty acid oxidation via CPT1b inhibition^84^. The acylcarnitine profile we demonstrate is consistent with peroxisomal dysfunction. Peroxisomes also generate a large class of plasmalogen phospholipid species ^85^. Plasmalogens have demonstrated anti-inflammatory effects and are protective against membrane ferroptosis^86^. In the current study, we show broad reductions in skeletal muscle plasmalogens consistent with reduced peroxisomal activity (Figure 6J). The reductions in skeletal muscle expression of Agpat3 (rate-limiting, Figure 6C) abundance of PC-O species, where “O” refers to the ether link at the sn-1 position (Figure 6L-M), provide additional evidence of peroxisomal dysfunction. Inherited peroxisomal disorders such as Zellweger syndrome show large reductions in plasmalogens and PC-O species, confirming the role of peroxisomes in the endogenous phospholipid synthesis^87^. Our results demonstrate that peroxisomal overload in HFpEF promotes phospholipid remodeling by shifting toward the synthesis of specific acylcarnitines to buffer acyl-CoA at the expense of lipid synthesis.

### Limitations of the Study

Our study provides the most in-depth analysis of skeletal muscle metabolism in HFpEF to date. We used a robust rodent model that phenocopies numerous features of cardiometabolic HFpEF, including skeletal muscle dysfunction^88^. Using multiomics integration, we have identified specific metabolic pathways that promote skeletal muscle dysregulation in the setting of cardiometabolic HFpEF. Our work is not without some limitations. First, our metabolomics data provide only a snapshot of metabolism, and therefore, the precise regulation of substrate flux requires additional isotope tracer approaches. We focused our analysis on metabolic pathways, but other cell signaling and inflammatory pathways could potentially yield new mechanistic insight into HFpEF pathobiology. We only studied male ZSF1 obese rats, which limits our ability to determine if sexual dimorphism is a factor in skeletal muscle dysregulation in HFpEF. While our work may be considered descriptive, very little molecular analysis has been conducted on skeletal muscle metabolism in cardiometabolic HFpEF. Future work utilizing skeletal muscle samples from patients with well-phenotyped cardiometabolic HFpEF patients will be necessary to confirm our findings.

## Conclusion

Skeletal muscle dysfunction is a major factor in exercise intolerance and frailty in HFpEF, but no studies have integrated multiomics data to define the potential sites of dysfunction. Our results and data from other groups strongly suggest that the skeletal muscle simultaneously undergoes a unique disease progression independent of cardiac dysfunction in HFpEF. This disease most closely resembles a metabolic myopathy. Our analysis provides a roadmap for future studies that will target HFpEF myopathy for the treatment of exercise intolerance and frailty in patients with cardiometabolic HFpEF.

## Acknowledgments

The authors thank Dr. Jarek Staszkiewicz for assistance with bioinformatic analysis.

